# PTP-3(LAR PTPR) promotes intramolecular folding of SYD-2(liprin-α) to inactivate UNC-104(KIF1A) in neurons

**DOI:** 10.1101/723288

**Authors:** Muniesh Muthaiyan Shanmugam, Syed Nooruzuha Barmaver, Hsin-Yi Huang, Oliver Ingvar Wagner

## Abstract

This study aims to demonstrate how PTP-3 regulates SYD-2 to control UNC-104-mediated axonal transport. UNC-104 is the *C. elegans* homolog of kinesin-3 KIF-1A known for its fast shuttling of STVs (synaptic vesicle protein transport vesicles) in axons. SYD-2 is the homolog of liprin-α in *C. elegans* known to directly regulate UNC-104 as well as being a substrate of LAR PTPR (leukocyte common antigen-related (LAR) protein tyrosine phosphatase (PTP) transmembrane receptor) with PTP-3 as the closest homolog in *C. elegans*. CoIP assays revealed increased interaction between UNC-104 and SYD-2 in lysates from *ptp-3* knockout worms. Intramolecular FRET analysis revealed that SYD-2 predominantly exists in an open conformation state in *ptp-3* mutants. These assays also revealed that non-phosphorylatable SYD-2 (Y741F) exists predominately in folded conformations while phosphomimicking SYD-2 (Y741E) exists predominantly in open conformations. In *ptp-3* mutants, SNB-1 cargo accumulates in soma while at the same time UNC-104 motors increasingly cluster along initial segments of axons. Interestingly, the unc-104 gene is downregulated in *ptp-3* mutants that might explain the vesicle retention phenotype. More strikingly, the few visibly moving motors and STVs were overly active in neurons of these mutants. We propose a model in which the lack of PTP-3 promotes increased open conformations of SYD-2 that in turn facilitates UNC-104/SYD-2 interactions boosting motor and STVs moving speeds.

## Introduction

The transport of vital neuronal factors such as nucleic acids, lipids, enzymes, and organelles (synaptic vesicles, mitochondria etc.) is essential for proper neuronal survival and functions. UNC-104 is the homolog of KIF1A in *C. elegans* known to be a fast anterograde transporter of STVs (synaptic vesicle protein transport vesicles) regulated by active zone protein SYD-2 (liprin-α) [1-3]. Heritable genetic defects in motor proteins and their adaptors are linked to various neurological diseases such as adult-onset neurodegenerative disease (AOND), Parkinson’s disease (PD), Alzheimer’s disease (AD), Charcot-Marie-Tooth disease CMT, tauopathies, and amyotrophic lateral sclerosis (ALS) [4-9]. Understanding how motor proteins are regulated will provide insights on how these diseases develop. The following mechanisms reflect few examples of how kinesins are regulated: 1) Intramolecular folding deactivates kinesins [10]. 2) Cargo binding activates kinesins [11]. 3) Physical pulling via opposing motor dynein activates kinesins [12]. 4) Binding of small adaptor proteins such as SYD-2 and LIN-2 activates kinesin [2, 3]. 5) Phosphorylation regulates kinesins [13]. 6) Post-translational modifications (PTM) of microtubules (MT) as well as microtubule-associated proteins (MAPs) regulate kinesins [14, 15]. 7) Differential organization of MT in subcellular regions regulate kinesin directionality [16]. In this study, we will focus on how SYD-2 regulates UNC-104 via intramolecular folding coordinated by PTP-3 phosphatase. A broad set of kinases such as GSK3β, MAPKs, ERK1/2, p38 MAPKs, JNKs, Cdk5, Akt, PKA, CK2, PKC, and PINK1 regulate axonal transport either directly (affecting motors) or indirectly (via adaptor proteins) [13]. On the other hand, how kinesins are regulated by phosphatases, remains largely unknown. One example is that kinesin-like protein Eg5 and MKlp2 are substrates of PTEN and PP2A phosphatases to function in cell division [17-20].

PTP-3 is an ortholog of human LAR PTPRD (leukocyte common antigen-related protein tyrosine phosphatase transmembrane receptor type D) and LAR PTPRF (leukocyte common antigen-related protein tyrosine phosphatase transmembrane receptor type F). It is composed of three N-terminal Ig domains, eight FN3 domains, a transmembrane domain, and two C-terminal PTP domains D1 and D2 (with D2 possessing very little or null phosphatase catalytic activity) [21]. Northern blot analysis identified two isoforms of PTP-3 that express broadly in various tissues at early worm stages, but is restricted to the nervous system in the adult worm. Worms carrying a *ptp-3* mutation have the posterior body bulged at the larval stage L1 based on defects in epidermal morphogenesis [22]. It has been also demonstrated that the PTP-3A isoform is localized to synaptic regions along with other active zone proteins such as UNC-10, however, the PTP-3B isoform is localized to extrasynaptic regions. Null mutations of ptp-3 lead to synaptic defects as well as defects in axon guidance and neuroblast migration, while overexpression of the PTP-3B isoform can rescue axon guidance and neuroblast migration defects, but not synaptic defects [23]. LIP.1 (LAR-interacting protein 1, later renamed to liprin-α) functionally interacts with the D2 domain of LAR and is often localized at focal adhesion. Functional interactions between liprins and LAR have been known for more than two decades [24, 25], however, only recently a proteomic study had revealed that a tyrosine at position 741 in SYD-2 may act as a potential target for PTP-3 [26]. In *C. elegans*, a *syd-2* mutation *ju37* was found to cause diffused localization of synaptic markers such as SNB-1, syntaxin, and RAB-3. Other phenotypes caused by *syd-2* mutations (such as *ju37* or *ok217*) include reduced locomotion, defective egg laying as well as egg retention and hatch-bag phenotypes [27, 28]. We have previously shown that the physiological function of SYD-2 is regulated by an intramolecular folding via an interaction between the N-terminal CC (coiled coil) domains and C-terminal SAM (sterile alpha motif) domains [2, 29]. In *C. elegans*, SYD-2 acts primarily as a scaffolding protein to regulate synapse formation, stability, and organization along with other active zone proteins such as ELKS-1, RYS-1, UNC-10 and sentryn [30-37]. Apart from this function, we have previously demonstrated that SYD-2 acts as an adaptor protein to regulate UNC-104’s motility by interacting with the motor’s FHA domain through its SAM domains. This interaction likely stabilizes the motor’s dimeric state thus enhancing motor motility. Based on this knowledge, we hypothesize that extrasynaptical PTP-3B may be an important factor to regulate UNC-104 motor activity via modulating the intramolecular folding of SYD-2 (likely by dephosphorylating tyrosine 741).

## Results

### Functional and genetic interactions among UNC-104, SYD-2 and PTP-3

To understand if genetic interactions among the proposed PTP-3/SYD-2/UNC-104 pathway exist, we measured mRNA levels of both syd-2 and unc-104 genes in two different *ptp-3* mutant strains *ptp-3(ok244)* and *ptp-3(mu256)*. As mentioned in the Methods section, the *ok244* allele encodes for a frameshift and premature termination in the ptp-3 gene resulting in a lack of the PTP-3A isoform while the PTP-3B isoform is still expressed. On the other hand, the *mu256* allele has been characterized to be a loss-of-function mutation and worms do not express both isoforms [23]. The same study revealed that PTP-3A primarily locates at synapses while PTP-3B localizes at extrasynaptic sites. From Figure 1A it is evident that worms carrying the *ok244* allele do not reveal significant differences in syd-2 and unc-104 mRNA expression while in *mu256* mutants both syd-2 and unc-104 genes are significantly downregulated (Fig. 1D). These results may point to a genetic relation not only between ptp-3 and syd-2 (as expected) but also, unexpectedly, between ptp-3 and unc-104. To understand whether PTP-3 plays a role in reported functional interactions between UNC-104 and SYD-2 [2, 38], we performed co-immunoprecipitation assays. Interestingly, UNC-104/SYD-2 interactions are pronounced in *ptp-3(mu256)* mutants (Fig. 1E+F; with pulled-down UNC-104::GFP used for normalization). Since the amount of co-precipitated SYD-2 in *ptp-3(ok244)* mutants varied greatly (due to unknown reasons), we cannot conclude whether this mutant affects UNC-104/SYD-2 interactions (Fig. 1B+C).

**Figure 1:**
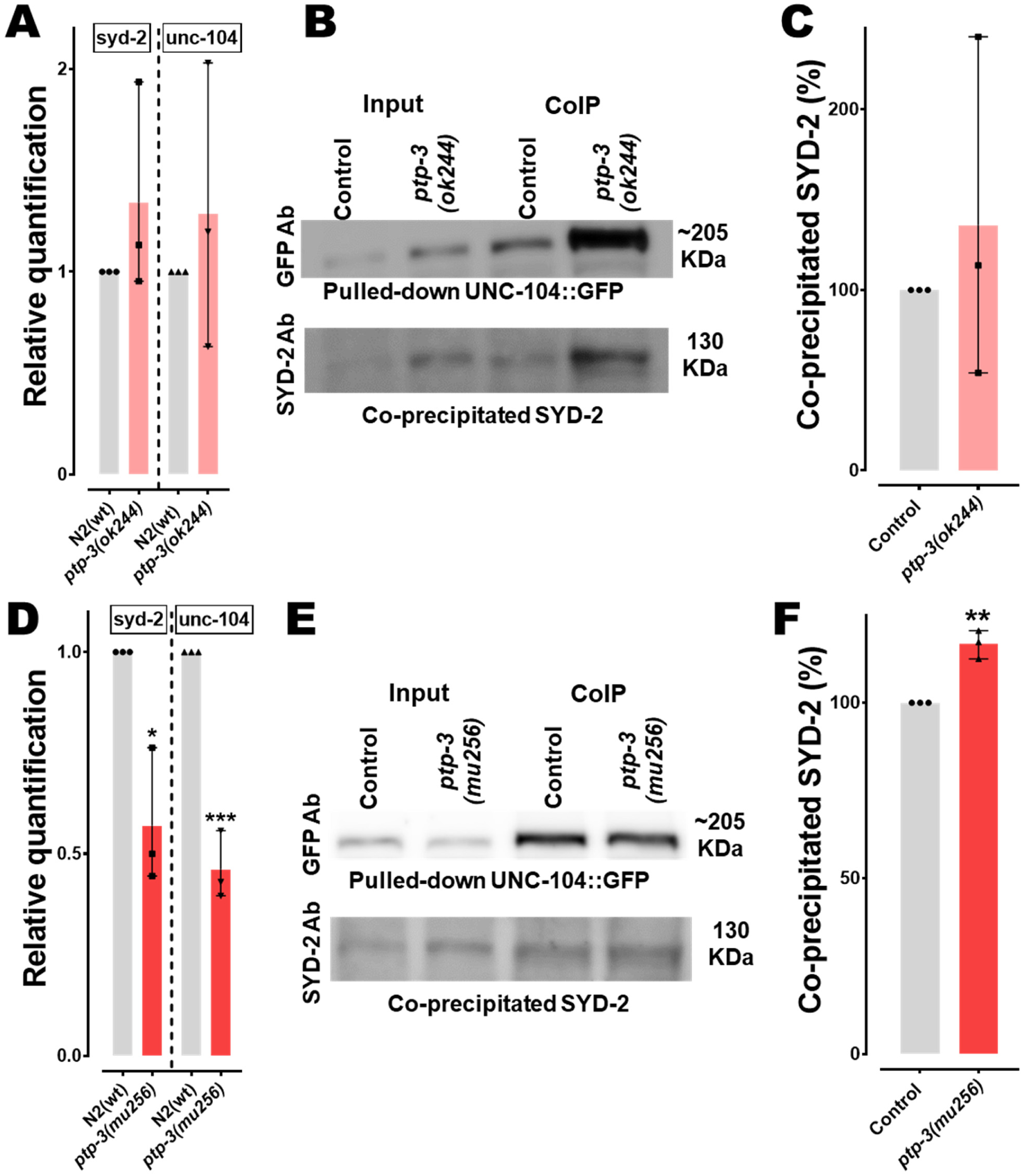
Changes in syd-2/unc-104 gene expression levels and SYD-2/UNC-104 protein interactions in *ptp-3* mutants. (A+D) Relative quantification of mRNA expression levels of syd-2 and unc-104 from lysates of young adult worms (N2 control, *ptp-3(ok244)* and *ptp-3(mu256)* allelic mutants). Quantitative PCR was performed thrice and each trial was performed in triplicates. (B+E) Upper lane: Western blot of pulled-down UNC-104::GFP using anti-GFP antibody. Lower lane: Blot of co-immunoprecipitated SYD-2 detected using anti-SYD-2 polyclonal antibody from lysates of *ok244* and *mu256* mutants. (C+F) Quantification of co-precipitated SYD-2 (normalized with UNC-104::GFP) from different genetic backgrounds. Error bars: ± max. and min. range. Unpaired Student’s t-test. * p<0.05, **p<0.01, ***p<0.001. t, df = 1.126, 4 (A, syd-2); 0.7039, 4 (A, unc-104); 4.389, 4 (D, syd-2); 11, 4 (D, unc-104); 0.6547, 4 (C) and 7.182, 4 (F).

To examine whether extrasynaptic PTP-3B isoform would function to regulate SYD-2, and thereby affect UNC-104 motor motility, we performed both colocalization as well as BiFC (bimolecular fluorescent complementation) assays [38]. Indeed, we revealed positive ICQ values when quantifying PTP-3B and SYD-2 colocalization in the *C. elegans* nervous system (Fig. 2 A-F; nerve ring and ventral nerve cord, VNC) as well as positive BiFC signals (Fig. 2G-J; nerve ring and VNC and ALM neuron). Critically, the PTP-3B/SYD-2 BiFC pair also colocalizes with a synaptic vesicle marker SNB-1 in the nerve ring (Fig. 2H+I). These data reveal that PTP-3 and SYD-2 have the potential for physical and functional interactions in the nervous system of *C. elegans*, not only at synapses (as reported by others for PTP-3A/SYD-2 [23]), but also within axons supporting our hypothesis for a potential role of PTP-3B/SYD-2 in UNC-104-mediated axonal trafficking.

**Figure 2:**
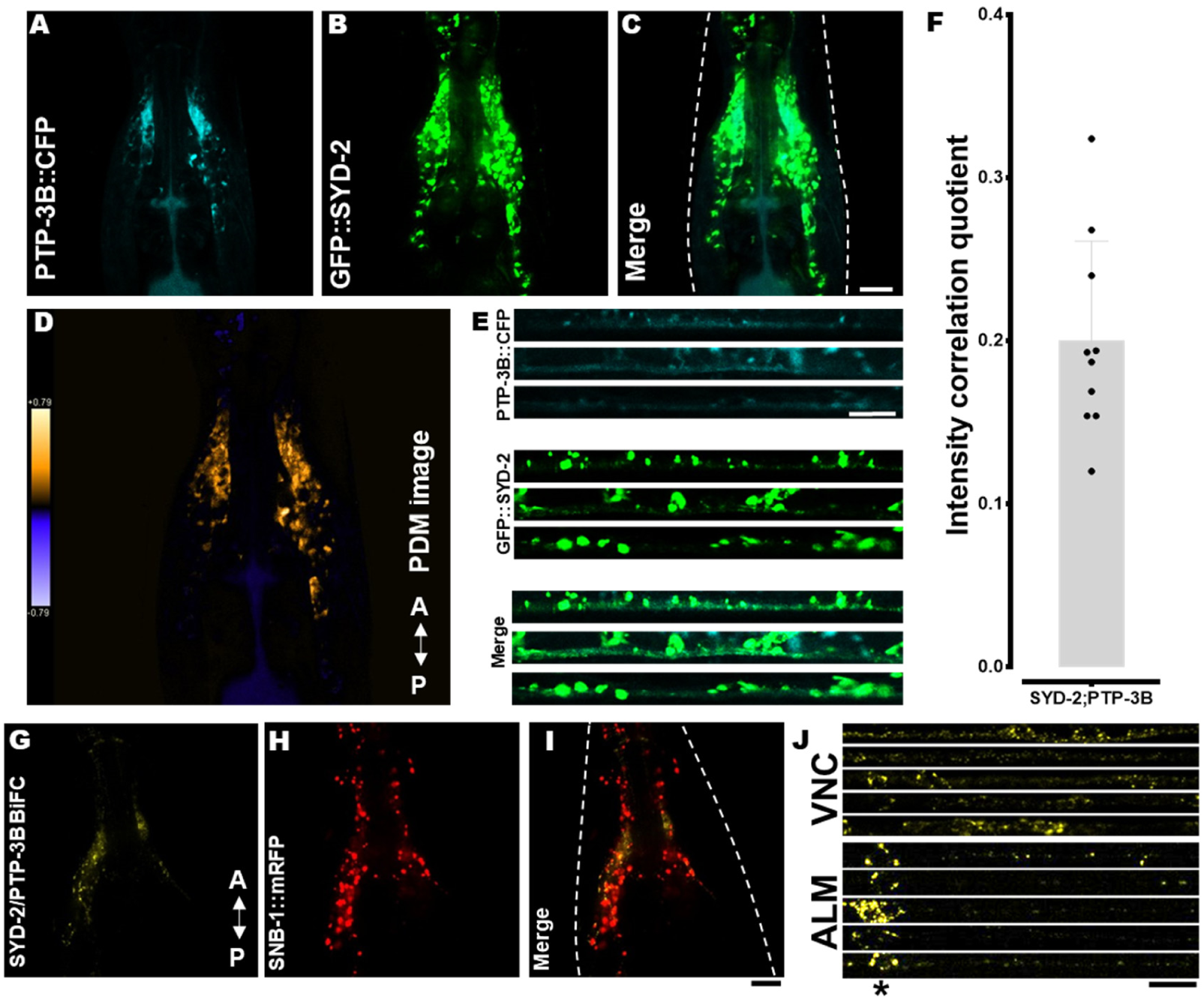
SYD-2/PTP-3B colocalization and BiFC (bimolecular fluorescence complementation) analysis. (A-E) PTP-3B::CFP and GFP::SYD-2 colocalization in nerve rings (A-D) and ventral nerve cords (E). (D) Pseudo-color PDM image (see Materials and Methods) indicating quantifiable colocalization between SYD-2 and PTP-3B in the nerve ring. (F) ICQ quantification of data shown in (C). (G) Positive PTP-3B/SYD-2 BiFC signal in the nerve ring of living worms indicating that SYD-2 and PTP-3B are at least 7-10 nm close to each other. (H) Worm expressing SNB-1::mRFP under a pan-neuronal promoter. (I) Merged image from (G) and (H). (J) Stacks of straightened VNCs (ventral nerve cords) and ALM neurons with positive PTP-3B/SYD-2 BiFC signals. * – indicates the location of somas in ALM neurons. Scale bars: 10 µm. N = 10 worms in (F). Error bar: ± max. and min. range. A – anterior direction and P – posterior direction.

### Lack of PTP-3B triggers increased unfolded states of SYD-2

Because SYD-2 exists in functional folded states, we propose that intramolecular folding of SYD-2 affect its binding propensities to UNC-104. As mentioned above, we assume that in *ptp-3* mutants diminished dephosphorylation of SYD-2 at Y741 would lead to increased unfolded states that would enhance its interactions with UNC-104. To approach this question we employed intramolecular FRET assays[39] as depicted in Figure 3A. From Figure 3B and C it is evident that the FRET ratio (IFRET/ID) of the Cypet::SYD-2::Ypet fusion is significantly reduced in *ptp-3(mu256)* mutants, indicating that unfolded states of SYD-2 are more pronounced in the absence of PTP-3. Critically, also knocking down the ptp-3 gene using RNAi led to comparable observations (Fig. 3C; Suppl. Fig. 3). As a control, we designed a non-phosphorylatable SYD-2 mutant Y741F and its FRET ratios are comparable to that of Cypet::SYD-2::Ypet expressed in N2 wild type animals. Moreover, a phosphomimicking SYD-2 Y741E construct led to FRET ratios comparable to that of the unmodified Cypet::SYD-2::Ypet construct expressed in *ptp-3(mu256)* mutants (Figs. 3B+C). These findings demonstrate that SYD-2 appears in a more pronounced open conformation in the absence of PTP-3 and that this mechanism is likely a function of the phosphatase activity of PTP-3 on tyrosine 741 of SYD-2.

**Figure 3:**
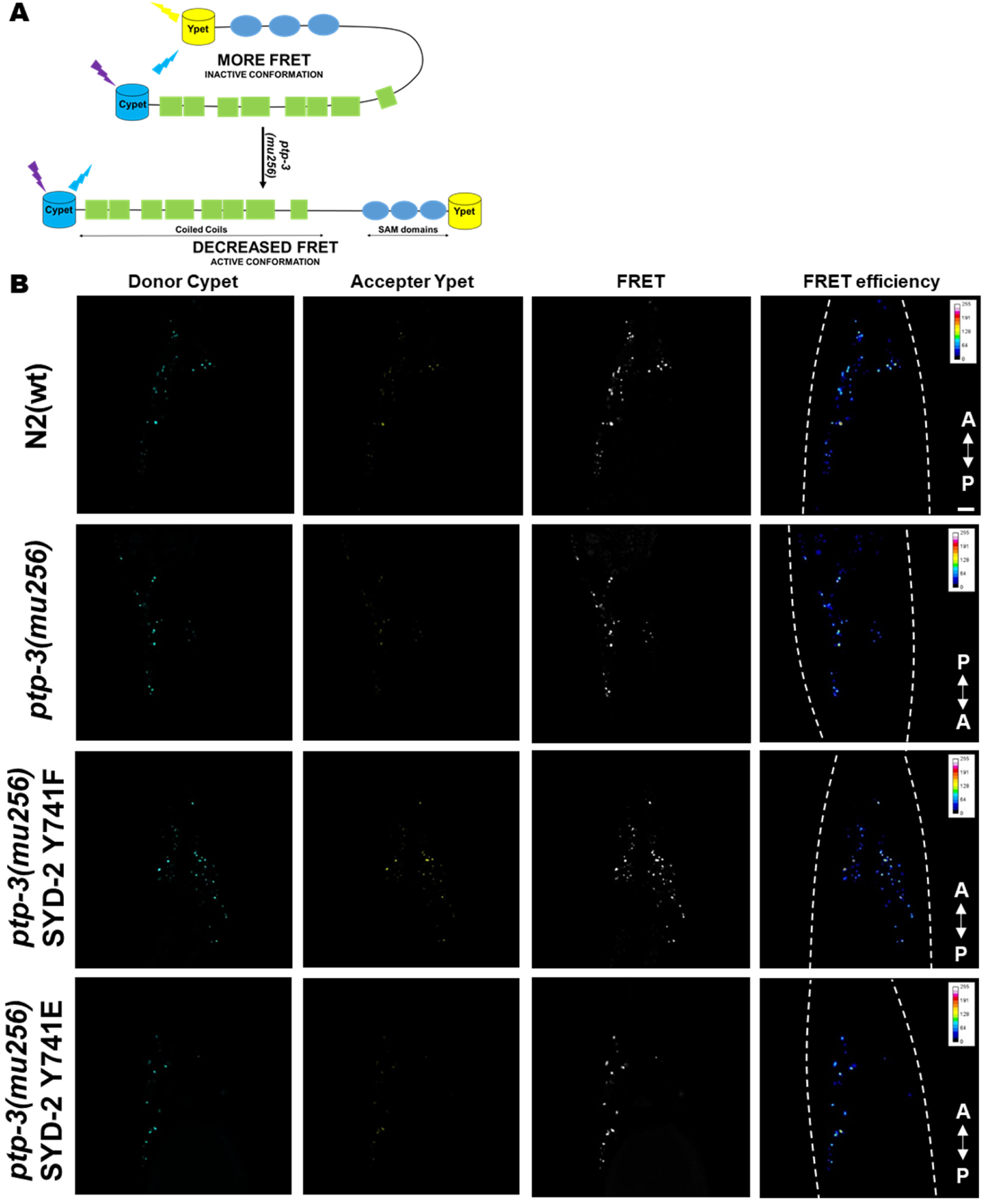
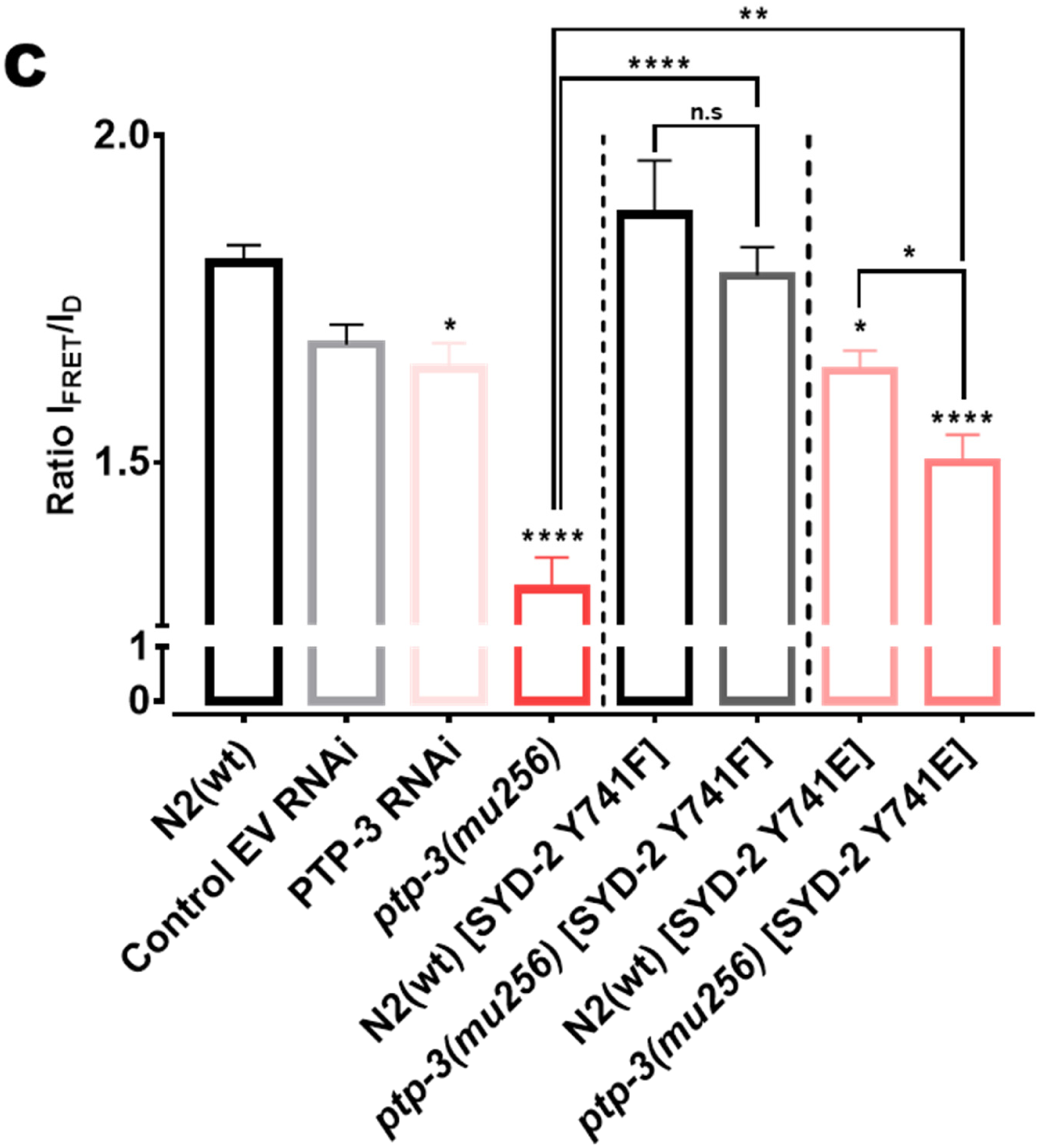
SYD-2 exists predominantly in an open conformation in *ptp-3(mu256)* mutants. (A) Pictorial representation of the designed intramolecular FRET assay for this study. (B) Images of *C. elegans* nerve rings from the respective fluorescence channel. (C) Quantification of FRET efficiencies of Cypet::SYD-2::Ypet in different genetic backgrounds, where ptp-3 RNAi, *ptp-3(mu256)* and phosphomimicking SYD-2 Y741E all lead to significant decrease in FRET ratios while the non-phosphorylatable SYD-2 Y741F reveals wild type signals. Scale bar: 10 µm. N = 26 images (C). Error bars: ± max. and min. range. One-way ANOVA with Fisher’s LSD test. * p<0.05, **p<0.01 and ****p<0.0001. F (DFn, DFd) = 15.83 (7, 200). A – anterior direction and P – posterior direction.

### Abnormal sorting of STVs and UNC-104 motor distribution in *ptp-3(mu256)* mutants

To understand whether this mechanism has any physiological significance, we analyzed STV distributions (with SNB-1 as a marker) in neurons in the presence and absence of PTP3B (Fig. 4A). SNB-1 particle sizes (Fig. 4B) are significantly reduced in the IAR (initial axonal region) of *mu256* mutant axons, while SNB-1 cargo density (Fig. 4C) as well as travel distances (Fig. 4D) remained unchanged in these mutants. Interestingly, we observed SNB-1 particle retention phenotypes in somas of HSN and ALM neurons (Fig. 4E-G). While cargo obviously accumulates in the soma of HSN neurons, it is at the same time reduced at synapses in these neurons (Fig. 4E+G; note that synapses of ALM neurons are not easily accessible for quantification). Because unc-104 gene expression is reduced in *mu256* mutants (Fig. 1D), we may explain the cargo retention phenotype partially with that finding.

**Figure 4:**
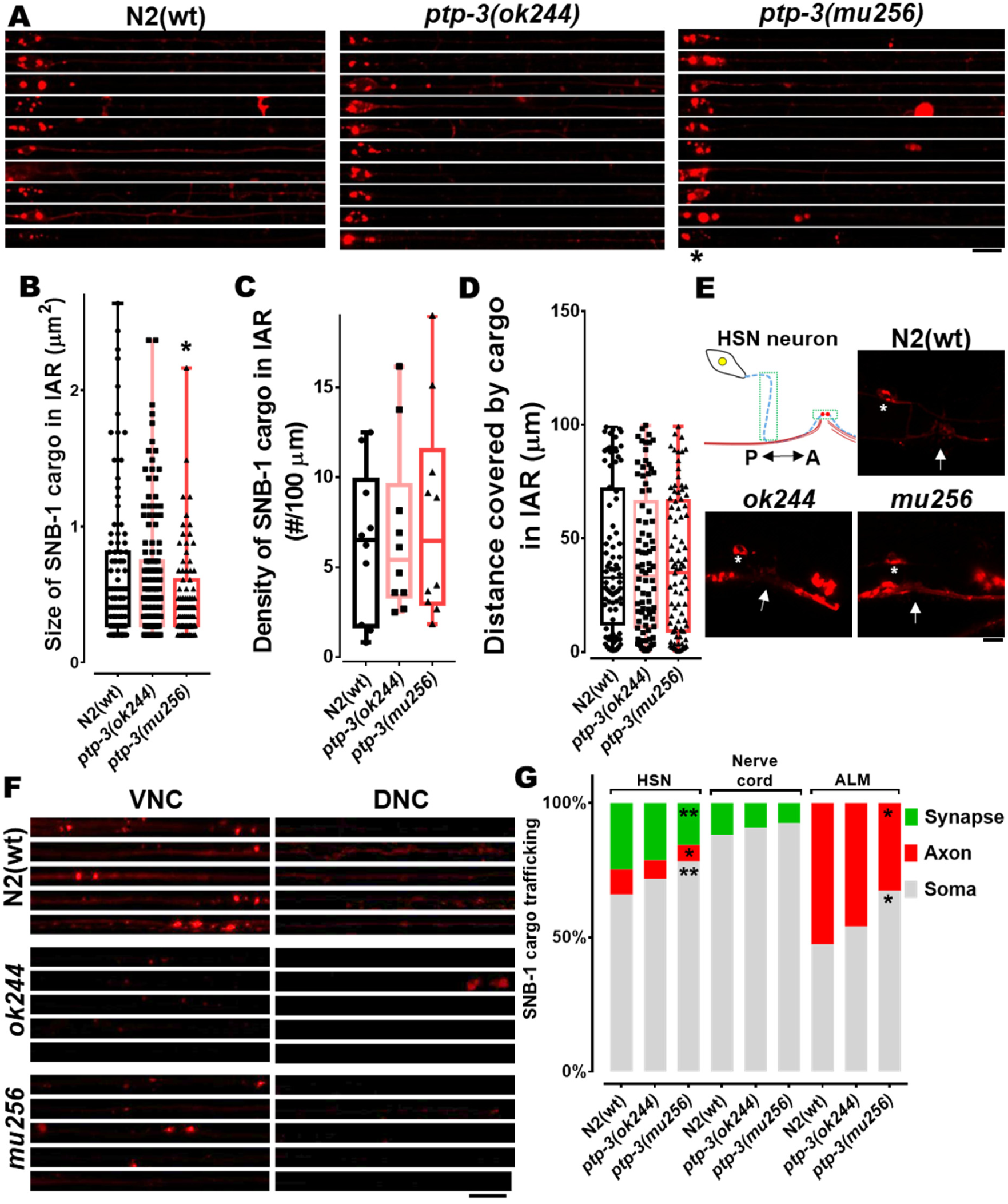
STVs accumulate in somas of *ptp-3* mutants. (A) Straightened ALM neurons taken from different worms within a population and stacked upon each other. (B-D) Quantification of size (B), densities (C) and travel distances (D) of SNB-1 containing synaptic vesicles from images displayed in (A). (E) Cartoon of HSN neuron with its presynaptic area and representative images of SNB-1 cargo distribution in wild type and mutants. * – indicates soma of neuron. White arrow indicates the location of the vulva. Green boxes indicate the region used for quantification of fluorescence intensity of axons (blue dashed line) and synapses (red circles). P – posterior direction and A – anterior direction. (F) Representative images of VNC and DNC from different study groups, straightened and stacked. (G) Quantification (fluorescence intensity) of the amount of cargo in the cell body and the initial axonal region of ALM, nerve cord (VNC-somas and DNC-synapses) and HSN neurons (soma, axon and synapse). Scale bars: 10 µm. Box plot with median and error bars: ± max. and min. range. One-way ANOVA with Fisher’s LSD multiple comparison test. * p<0.05 and ** p<0.01. F (DFn, DFd) = 3.426 (2, 320) (B); 3.862 (2, 27) (G, ALM); 4.327 (2, 43) (G, HSN soma); 2.455 (2, 43) (G, HSN axon) and 3.909 (2, 43) (G, HSN synapse).

We then analyzed the distribution patterns of UNC-104 motor in the long axons of sub-lateral neurons in *ptp-3* and *syd-2* mutants as well as in *ptp-3*;*syd-2* double mutants. As a result, density of UNC-104 motor clusters increased significantly in *ptp-3(mu256)* mutants compared to wild types, and this phenotype can be rescued when overexpressing PTP-3B in *mu256* mutants (Figs. 5A+C). Critically, UNC-104 motor clustering was reduced in *syd-2* knockout animals (Fig. 4C) consistent with previous findings [2]. As expected, overexpression of non-phosphorylatable SYD-2 Y741F was unable to rescue the *ok217* effect, meaning the (expectedly) increased closed conformations of this SYD-2 version constrained interactions with UNC-104. On the other hand, the phosphomimicking version of SYD-2 led to significantly increased UNC-104 clustering (Fig. 5C), probably due to increased UNC-104 scaffolding along axons via SYD-2 [2]. Epistasis analysis employing *syd-2;ptp-3* double mutants revealed that lack of PTP-3 does not affect the *syd-2* phenotype (Figs. 5A+C), thus, PTP-3 is likely located upstream of SYD-2.

**Figure 5:**
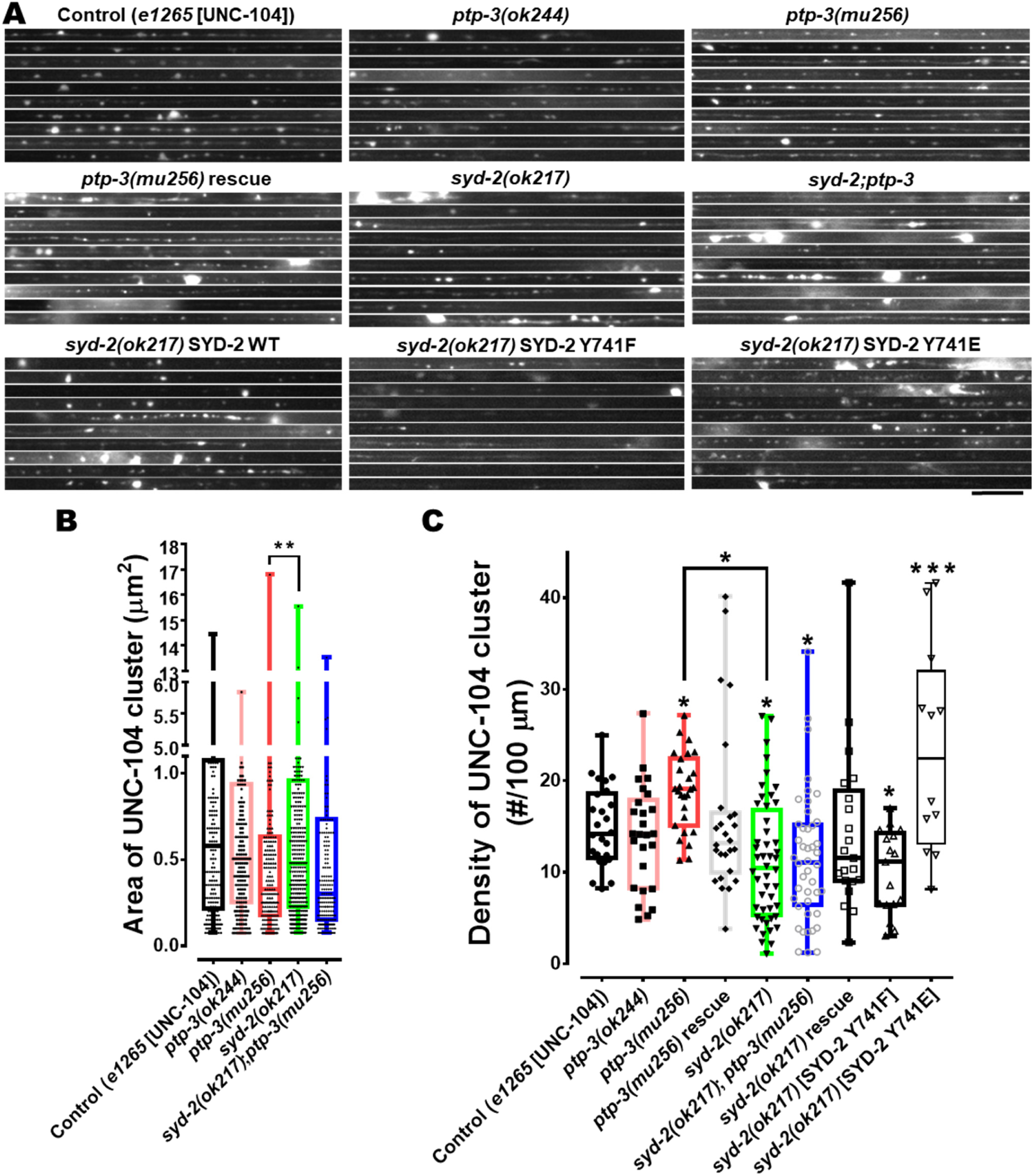
UNC-104 clustering is increased in *ptp-3* mutants and decreased in *syd-2* mutants. (A) UNC-104 accumulations and distribution patterns in straightened segments of sub-lateral neurons taken from different worms within a population and stacked upon each other. (B) UNC-104 cluster sizes in sub-lateral neurons. (C) UNC-104 densities along sub-lateral neurons. Scale bar: 10 µm. Box plot with median and error bars: ± max. and min. range. One-way ANOVA with Fisher’s LSD multiple comparison test. **** p<0.0001, ** p<0.005 and * p<0.05. F (DFn, DFd) = 7.073 (8, 238) (C).

### PTP-3 knockout results in increased anterograde motor and cargo velocities

Based on the idea that PTP-3 suppresses the function of SYD-2 (that in turn would deactivate UNC-104), it is crucial to analyze UNC-104’s motility behavior in *ptp-3* mutants. Indeed, kymograph analysis revealed that UNC-104 velocities in *ptp-3(mu256)* mutants significantly increased in anterograde directions (Fig. 6A; see also Suppl. Fig. 5+6). Because UNC-104 is a plus-end directed motor, it is not surprising that we did not measure significant changes in retrograde directions (Fig. 6B). However, it needs to be noted that UNC-104 may exist in complexes with dynein [12] or that multiple motors of different directionalities (antagonistic motors) may accumulate and bind to the same cargo vesicle resulting in saltatory movements of that vesicle. Because we only observe the movements of fluorescent UNC-104 particles, we may expect that we measure either moving parameters of single motors or those of multiple motors accumulated on a single vesicle. Nevertheless, it is obvious that bidirectional movements of UNC-104 in *ptp-3(mu256)* mutants increased (Fig. 6C) pointing to increased tug-of-war events (such as between UNC-104 and dynein). Critically, the observed increase in anterograde velocities of UNC-104 in *ptp-3(mu256)* mutants could be easily rescued by overexpressing a full-length PTP-3B in *mu256* mutants (Fig. 6A). Noteworthy, no significant changes of UNC-104 moving properties in *ptp-3(ok244)* mutants were detected, likely as a result of unaffected isoform PTP-3B in these mutants (Suppl. Fig. 1).

**Figure 6:**
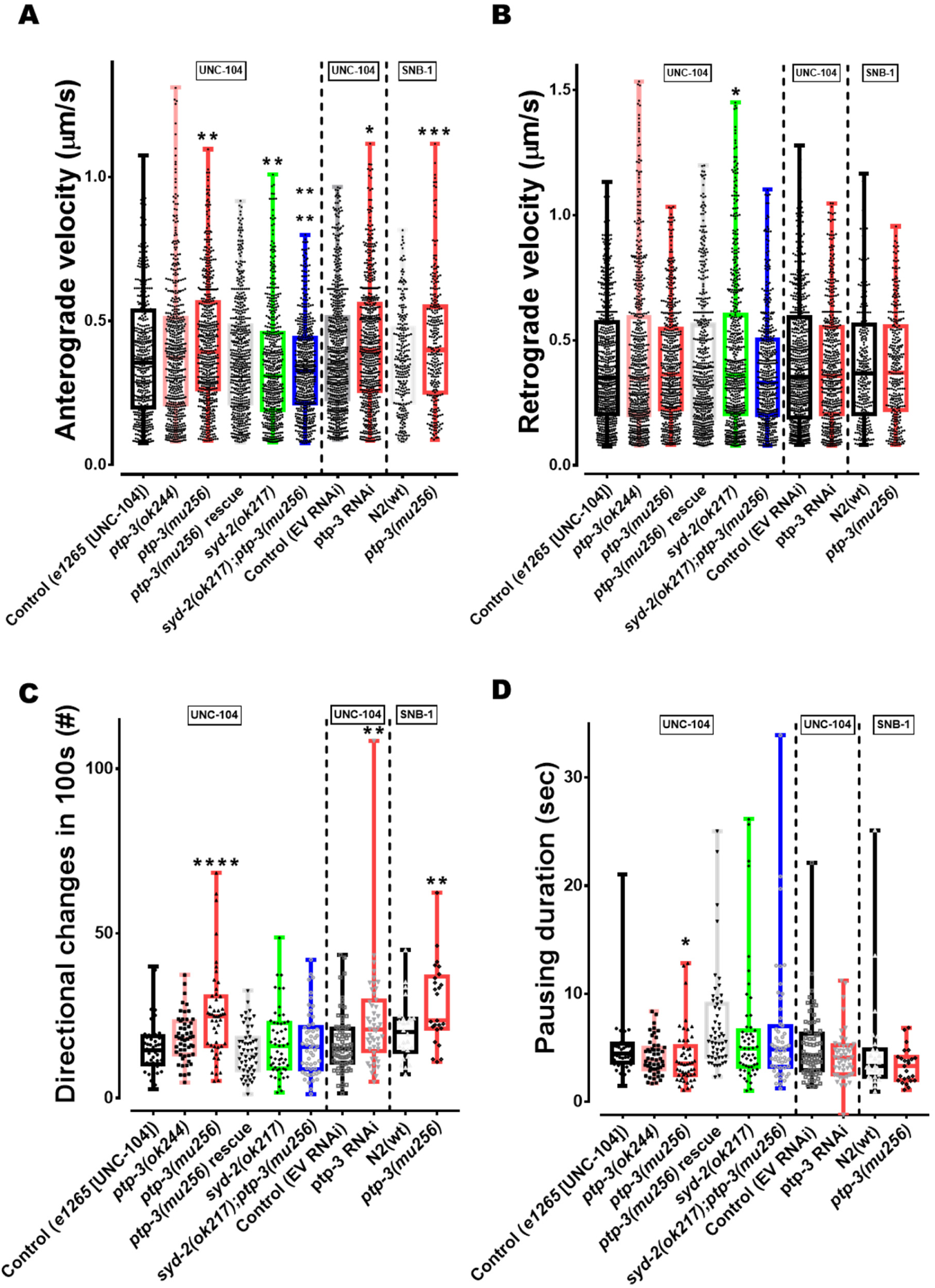
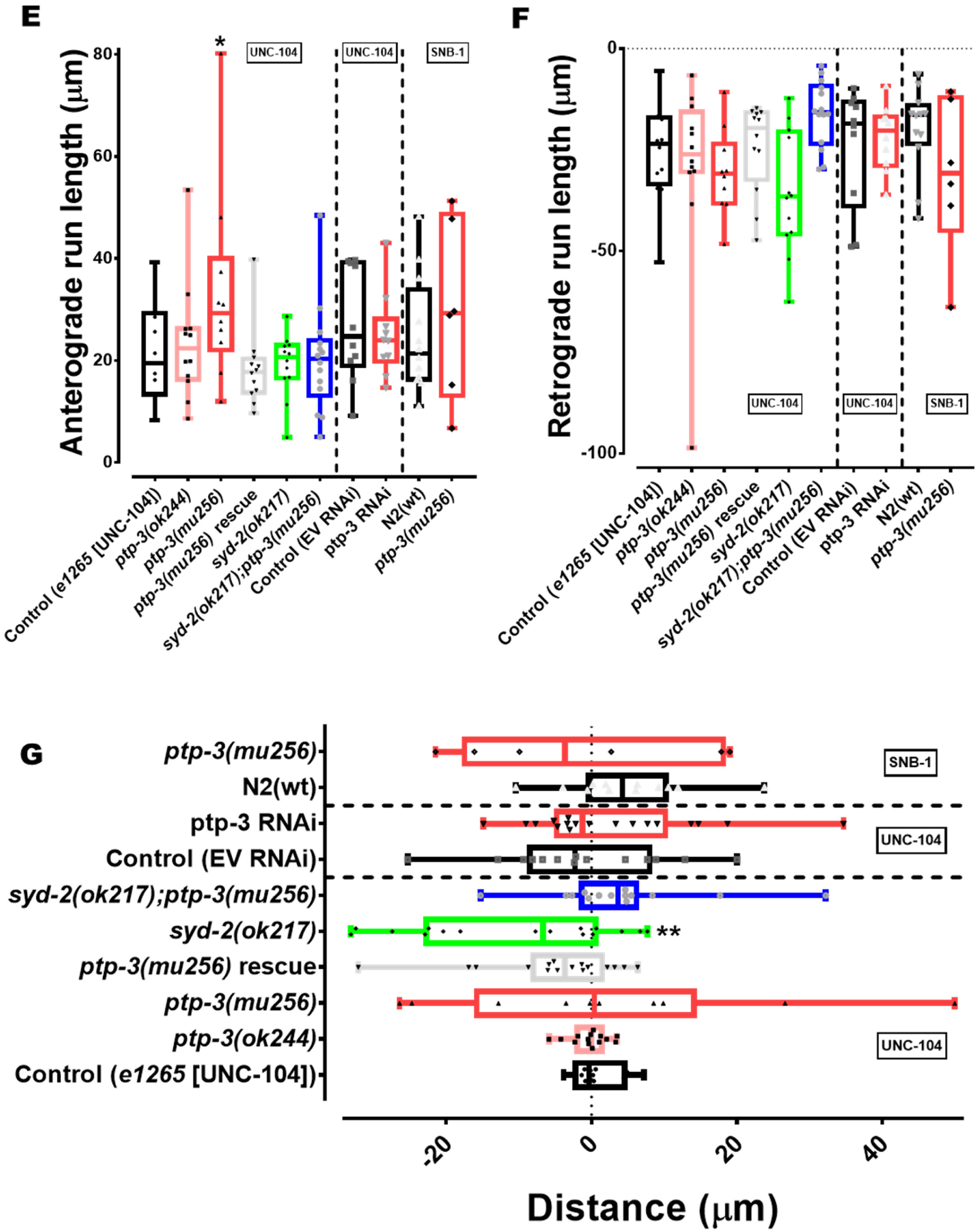
Both motor and cargo speeds are increased in *ptp-3* mutants. (A) UNC-104 and SNB-1 anterograde velocities in various genetic backgrounds as well as after ptp-3 RNAi. (B) UNC-104 and SNB-1 retrograde velocities in various genetic backgrounds as well as after ptp-3 RNAi. (C) Quantification of motor and cargo reversals (directional changes). (D) Quantification of motor and cargo pausing duration. (E+F) Quantification of total anterograde (E) and (F) retrograde run length. (G) Quantification of net run length. Analyzed UNC-104 particles: > 1500 events, SNB-1 particles: >750 events. Box plots with median and error bars: ± max. and min. range. One-way ANOVA with Fisher’s LSD multiple comparison test for UNC-104 in mutant strains and t-test with Welch’s correction for RNAi and SNB-1. **** p<0.0001, *** p< 0.001, ** p<0.005, and * p<0.05. F (DFn, DFd) = 12.65 (5, 3210) (A); 4.257 (5, 3394) (B); 8.452 (5, 309) (C); 4.614 (5, 310) (D); 2.515 (5, 66) (E); 2.778 (5, 66) (F) and 3.324 (5, 78) (G).

As reported earlier, UNC-104 anterograde velocities are significantly reduced in *syd-2(ok217)* knockout animals [2] and we reproduced this finding (Fig. 6A). Moreover, this effect cannot be rescued by the *ptp-3* mutation, once more confirming that PTP-3 is likely located upstream of SYD-2 in a single pathway. Importantly, moving characteristics of UNC-104’s major cargo SNB-1 are very consistent with that observed for the motor alone: both anterograde velocities as well as tug-of-war (reflected by particle directional changes) of SNB-1 particles are increased in *mu256* mutants (Figs. 6A+C). These observations emphasize the consistency of measurements and stresses the overall conclusion that the presence of PTP-3 would act to reduce axonal trafficking via the inhibition of SYD-2’s activating function on UNC-104. Besides investigating these effects in knockout worms, RNAi knockdown of the ptp-3 gene in N2 worms lead to comparable results (Figs. 6A-D). Besides these parameters (motor velocities and reversals) it is critical to note that anterograde total run length of motors increased in PTP-3 knockout animals (Fig. 6E); however, no differences in retrograde total run length (Fig. 6F) as well as in net run length (Fig. 6G) were detected. These results are consistent with the unaffected travel distances of STVs in *ptp-3* mutants (Fig. 4D).

## Discussion

### Identification of a novel pathway that regulate anterograde axonal transport

The role of SYD-2 as an UNC-104 activator, its potential to undergo intramolecular folding as well as its amino acid Y741 as a potential target for PTP-3 dephosphorylation have been reported previously [2, 26, 29]. From these studies, we hypothesized that PTP-3 may act as a regulator of SYD-2 to control kinesin-3 motor activity and above we provided a large set of evidences that support our hypothesis. Using SILAC–based quantitative phosphoproteomics, Mitchell *et al.* 2016 demonstrated that SYD-2 is a substrate of PTP-3, and that in *ptp-3* mutants SYD-2’s amino acid Y741 (located near the central stalk domain, Suppl. Fig. 1) is 40% hyperphosphorylated [26]. Although, it is a common practice to employ alanine and glutamic acid to design non-phosphorylatable and phosphomimicking mutants, one study has demonstrated that phenylalanine is a more appropriate substitution to create non-phosphorylatable proteins. The same study implied that glutamic acid substitution for tyrosine is insufficient to reproducibly gain the desired phosphomimicking effect [40]. However, we have clearly demonstrated that Y/E phosphomimicking is largely comparable to wild type hyperphosphorylated SYD-2 in *ptp-3(mu256)* mutant backgrounds (Fig. 3C). Combined with the co-immunoprecipitation data (Fig. 1), it can be inferred that the open conformation of SYD-2 leads to exposed SAM domains (necessary for UNC-104 binding [2]) resulting in elevated interactions with UNC-104 leading to increased motor activation.

Noteworthy is also that other regulators such as RSY-2 may regulate SYD-2 and it is known that this protein acts to suppress SYD-2-dependent synapse assembly, specifically because of unexposed N- and C-terminal domains of SYD-2 incapable to scaffold with other active zone proteins [29, 41]. Also, Chia *et al.* 2013 concluded that although only CC (coiled coil) domains of SYD-2 are sufficient to assemble functional synapses, a L1086F mutation in the SAM domain of full-length SYD-2 results in loss of synapses, probably due to increased folding of SYD-2 [29]. Several proteins are known to interact with either only SYD-2’s CC domains (ELKS-1, UNC-10 and GIT-1) or only SYD-2’s SAM domains (PTP-3 and GRIP) [23, 31, 42, 43]. Intramolecular folding of SYD-2 will likely mask such interaction sites preventing complex interactions with such proteins. Critically, proteins such as UNC-104 and LIN-2 have been shown to directly interact with both CC and SAM domains at functionally different degrees [2, 3].

Liprin-α has been shown to interact with ATP-agarose beads leading to the assumption that it may possess the ability of autophosphorylation [44], however, there is no known kinase-domain in liprins needed for such regulation. Stronger evidence for another SYD-2 regulator is CASK/LIN-2 that has been shown to cooperatively act (together with SYD-2) to regulate UNC-104 motility, specifically on the motor’s superprocessivity [3]. Also, it has been shown that a calcium/calmodulin-dependent protein kinase II colocalizes with liprin-α to regulate the degradation of liprin-α via phosphorylation [45]. Lastly, a recent study shows that UNC-16, a MAP-kinase like protein regulates the CDK-5/SAD-1/SYD-2 triplet protein system to inhibit organelle trafficking in neurons [46]. Further studies are needed to unravel the type of kinase that phosphorylates amino acid Y741 in SYD-2.

### Physiological effects of the PTP-3/SYD-2/UNC-104 pathway

Because mRNA levels of syd-2 and unc-104 were significantly reduced in *mu256* mutants (Fig. 1D), it is likely that STV vesicle trafficking is compromised in these animals consistent with other studies [1, 36, 47-49]. Indeed, mild accumulation of STVs in somas of ALM and HSN neurons, as well as reduced STVs in axons or synapses, can be seen in PTP-3 knockout worms (Fig. 4G). While based on this finding it is obvious that *fewer* motors are available for axonal transport, yet, these motors are *more activated* (moving at faster speeds, Fig. 6A). This increased motor activation phenotype is likely based on an elevated number of activated SYD-2 molecules due to increased open states in *mu256* mutants (Fig. 3). One study in *Drosophila* Dlar mutants revealed that synaptic boutons are reduced as well as excitatory postsynaptic potential is decreased [50]. Another study in rat neurons demonstrated that both LAR RNAi and dominate-negative disruption of LAR results in reduced number of excitatory synapses as well as dendritic spines. The same study uncovered that LAR/liprin-α interaction is essential for dendritic targeting of AMPA receptors at postsynaptic terminals [51]. These findings may explain the observed decreased sizes of STVs in the axon of ALM neurons (Fig. 4B), and it is noteworthy that one study relates the size of STVs to the osmotic pressure (regulated by neurotransmitter concentration) within the STVs [52]. In *Drosophila*, the amount of neurotransmitter in vesicles increased along with STV size upon overexpression of vesicular glutamate transporter [53, 54]. All these findings lead to the assumption that reduced vesicle sizes (Fig. 4B) could be due to diminished amounts of neurotransmitters inside STVs and, moreover, abnormal STV sorting (Fig. 4G) may trigger reduced expression of syd-2 and unc-104 (Fig. 1D). A recent study in *Drosophila*, uncovered an activation of Wnd/DLK MAP kinase signaling after disruption of Unc-104’s function (as well as upon overexpression of presynaptic proteins), leading to restrained expression of active zone components (in a feedback circuit) [55]. Impeded synapse formation may also explain the mild egg retention phenotype in *ptp-3(mu256)* mutants (Supp. Fig. 2). Egg laying in *C. elegans* is regulated via HSN neurons that innervate vulval muscles. Indeed, proper formation of synapses via the interplay of SYG-1, SYD-1 and SYD-2 are essential for functional egg-laying, and in *syd-1* as well as *syd-2* mutants, observed egg-laying defects often result in hatch-bag phenotype (Supp. Fig. 2) [27, 28].

SYD-2 has been described to inherit two major roles on UNC-104: one is the motor scaffolding function and the second is motor activation. In N2 wild type animals UNC-104 appears clustered along axons and this clustering has been shown not to be related to overexpression effects, and further, the patterns of UNC-104 clustering are different from the regular (pearl string-like) pattern of *en passant* synapses [2, 14]. Based on this knowledge one can deduce that open SYD-2 states in *ptp-3* mutants (Fig. 3) may trigger increased UNC-104 clustering along axons. Indeed, motor accumulation analysis exposed that not only UNC-104 clustering along axons is reduced in *syd-2* mutants (Fig. 5A+B), but also that PTP-3 may act upstream of SYD-2 (based on double mutant analysis) (Fig. 5C). This finding is in accordance with investigations from *Drosophila* demonstrating that Dliprin-α is essential for Dlar dependent synapse assembly [50].

Because axonal distribution patterns of SNB-1 and UNC-104 in axons, as shown in Figures 4 and 5, only reflects a static snapshot of cargo displacements, dynamic motility analysis of cargo and motor is thus vital as well. Critically, motility data as shown in Fig. 6 go well along with the findings from cluster pattern analysis (Figs. 4+5) and both clearly reveal the suppressing role of PTP-3 on SYD-2 and its function in axonal transport (summarized in Fig. 7). Moreover, motility features of the motor *alone* do nearly perfectly mirror the motility features of the cargo *alone* (Fig. 6) underlining the validity of these observations. Worth discussing are also the elevated motor reversals in *ptp-3* mutants (Fig. 6C) pointing to increased tug-of-war events. Future studies need to investigate the role of dynein and it might be suspected that in dynein mutants motor reveals remain unaltered. Nevertheless, based on the “paradox of co-dependence” and “mechanical activation” theories [56], increased directional changes may be based on hyperactivated UNC-104 and thus mechanically activating dynein (if both bound to one vesicle). Though this might be valid for short-term runs, the determined considerable increase in total anterograde run lengths in *ptp-3* mutants (Fig. 6E) could be rather based on the motor coordination model for longer runs. Critically, these observations go well along with current views in the motor community suggesting that tug-of-war events are mostly observed for short runs [57].

**Figure 7:**
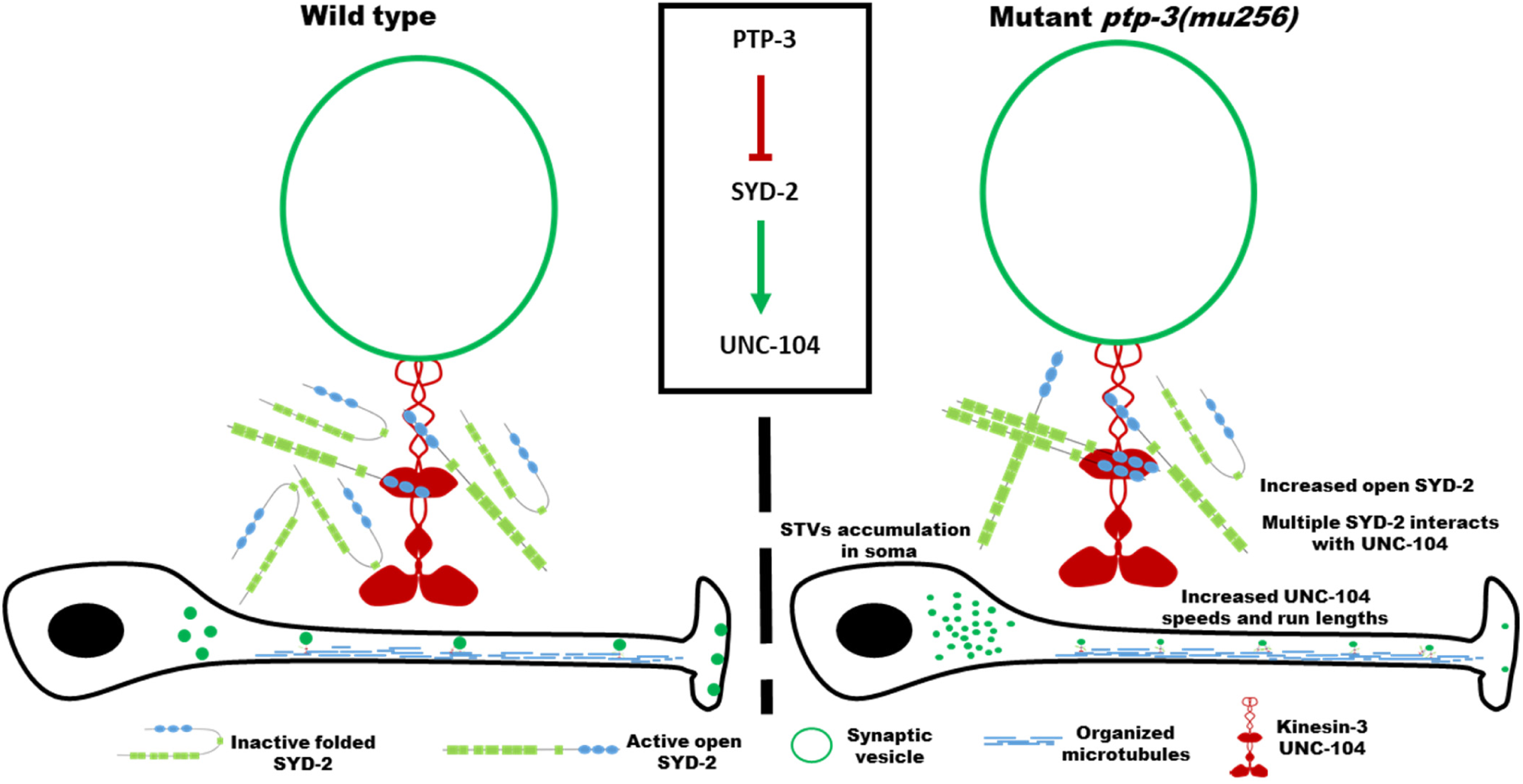
Model of PTP-3 to promote folded states of SYD-2 to inactivate UNC-104. Pictorial summary of findings from this study. PTP-3 is upstream of SYD-2 that regulates UNC-104 activity. PTP-3 dephosphorylates SYD-2 at position Y741 resulting in intramolecular folding of SYD-2 which concomitantly inactivates UNC-104. Vice versa, lack of PTP-3 results in more open configurations of SYD-2 resulting in increased interaction with UNC-104 [3] promoting increased UNC-104 scaffolding along the axon and activating UNC-104 with visibly increased speeds. However, STVs accumulate in the soma due to reduced expression of syd-2 and unc-104 in *ptp-3* mutants.

## Materials and methods

### Plasmid constructs and *C. elegans* strains

Worm strains were maintained and analyzed at 20°C on NGM (nematode growth medium) agar plates seeded with uracil auxotroph *E. coli* OP50 strain serving as nematode food source [58]. All analysis in this study were performed on young adult worms unless otherwise stated. Worm strains N2, RB633 *ptp-3(ok244)*, CZ3761 *ptp-3(mu256)*, ZM607 *syd-2(ok217)* were obtained from CGC (*Caenorhabditis* Genetic Center, MN, USA), and *unc-104(e1265)*;Ex[*Punc-104::unc-104::mrfp*], *unc-104(e1265)*;Ex[*Punc-104::unc-104::gfp*] and *syd-2(ok217)*;Ex[*Punc-104::unc-104::mrfp*] were described in [2, 3, 14]. Worm strains OIW 57 *ptp-3(ok244)*;nthEx57[*Punc-104::unc-104::mrfp*], OIW 58 *ptp-3(mu256)*;nthEx58[*Punc-104::unc-104::mrfp*], OIW 59 *ptp-3(ok244)*;nthEx59[*Punc-104::unc-104::gfp*] and OIW 60 *ptp-3(mu256)*;nthEx60[*Punc-104::unc-104::gfp*] were generated by crossing mutant strains with existing strain *unc-104(e1265)*;nthEx[*Punc-104::unc-104::mrfp*] or *unc-104(e1265)*;nthEx[*Punc-104::unc-104::gfp*]; double mutant OIW 61 *syd-2(ok217);ptp-3(mu256)*;nthEx61[*Punc-104::unc-104::mrfp*] was generated by crossing *syd-2(ok217)* mutant with males of OIW 58 *ptp-3(mu256)*;nthEx58[*Punc-104::unc-104::mrfp*]. Transgenic line OIW 62 *ptp-3(mu256)*;nthEx62[*Punc-104::unc-104::mrfp; Punc-104::ptp-3b::cfp*] was generated by microinjection of *Punc-104::unc-104::mrfp* [2] and *Punc-104::ptp-3b::cfp* (100 ng/µl each). Note that for the pan-neuronal Punc-104 promotor to fully operate, relatively high dosages of plasmids are necessary [59]. PTP-3B containing first 6 introns and exons followed by cDNA was cloned into already existing *Punc-104::cfp* plasmid (generated by replacing *gfp* in *pPD95.77* with *cfp* and subsequently cloned with *Punc-104*) by homologous recombination (In-Fusion^®^ HD cloning, Takara Bio USA, Inc.) using CTCTAGAGGATCCCCATGGGAACGCCGGCT (forward) and CCTTTGGCCAATCCCTTCGAGAAATTGTCATAGGCGGC (reverse) primers. Genomic DNA of snb-1 gene was amplified and cloned into *Punc-104::mrfp* using homolog recombination with CTCTAGAGGATCCCCATGGACGCTCAAGGAGATGCC (forward) and CCTTTGGCCAATCCCTTTTTTCCTCCAGCCCATAAAACGATGAT (reverse) primers to generate *Punc-104::snb-1::mrfp* plasmid, and microinjected at 80 ng/µl to generate OIW 63 *N2(wt)*;nthEx63[*Punc-104::snb-1::mrfp*], OIW 64 *ptp-3(ok244)*;nthEx64[*Punc-104::snb-1::mrfp*], OIW 65 *ptp-3(mu256)*;nthEx65[*Punc-104::snb-1::mrfp*] transgenic lines. For colocalization studies OIW 66 *ptp-3(mu256)*;nthEx66[*Punc-104::gfp::syd-2; Punc-104::ptp-3b::cfp*] worm was generated by injection of *Punc-104::gfp::syd-2* [3] and *Punc-104::ptp-3b::cfp* (both at 100 ng/µl) plasmids. Plasmid construct *ptp-3b::vc155* was developed by subcloning ptp-3b gene from *Punc-104::ptp-3b::cfp* into *Punc-104::vc155* and microinjected along with *Punc-104::vn173::syd-2* [38] and *Punc-104::snb-1::mrfp* to generate OIW 67 *ptp-3(mu256)*;nthEx67[*Punc-104::vn173::syd-2; Punc-104::ptp-3b::vc155; Punc-104::snb-1::mrfp*] (75 ng/µl, 75 ng/µl and 50 ng/µl, respectively). For FRET (Förster/Fluorescence resonance energy transfer) experiments, the syd-2 gene from an already existing plasmid construct *Punc-104::gfp::syd-2* [3] was subcloned into *Punc-104::cypet::ypet* using homologous recombination with GATCCCCGGGATTGGAAATGAGCTACAGCAATGGAAAC (forward) and CTTTGGGTCCTTTGGTGGTATATAAATGAAACTCGTAGGATTTTGC (reverse) primer set. The created plasmid was microinjected at a concentration of 100 ng/µl with *snb-1::mrfp* as coinjection marker to generate OIW 68 *N2(wt)*;nthEx68[*Punc-104::cypet::syd-2::ypet; Punc-104::snb-1::mrfp*], transgenic worm strain. SYD-2 Y741F mutation was generated by whole plasmid amplification of *Punc-104::gfp::syd-2* [3] using 5’ phosphorylated TCGGCAATCCGCAGTTTGTTG (forward) and 5’ phosphorylated AAATATCATACCGTCGGTCACCGG (reverse) primers resulting in *Punc-104::gfp::syd-2 Y741F*; from the resulting mutant construct, SYD-2 Y741F, was subcloned into *Punc-104::cypet::ypet* to generate *Punc-104::cypet::syd-2 Y741F::ypet*. SYD-2 Y741E mutation was generated by overlap extension PCR using: ATGAGCTACAGCAATGGAAACATAAATTG (forward) and GCGGATTGCCTTCAATATCATACCGTCGGTCACCG (reverse) for 5’ segment of syd-2 and CGGTATGATATTGAAGGCAATCCGCAGTTTG (forward) and CTAGGTATATAAATGAAACTCGTAGGATTTTGCT (reverse) for 3’ segment of syd-2; subsequently subcloned to generate *Punc-104::cypet::syd-2 Y741E::ypet* and *Punc-104::gfp::syd-2 Y741E*. Other FRET mutant strains OIW 69 *ptp-3(mu256)*;nthEx69[*Punc-104::cypet::syd-2::ypet; Punc-104::snb-1::mrfp*], OIW 70 *ptp-3(mu256)*;nthEx70[*Punc-104::cypet::syd-2Y741F::ypet; Punc-104::snb-1::mrfp*], OIW 71 *N2(wt)*;nthEx71*Punc-104::cypet::syd-2Y741F::ypet; Punc-104::snb-1::mrfp*], OIW 72 *ptp-3(mu256)*;nthEx72[*Punc-104::cypet::syd-2Y741E::ypet; Punc-104::snb-1::mrfp*] and OIW 73 *N2(wt)*;nthEx73[*Punc-104::cypet::syd-2Y741E::ypet; Punc-104::snb-1::mrfp*] were generated by microinjection of aforementioned FRET constructs at 120 ng/µl along with snb-1::mrfp coinjection marker. Rescue of *syd-2(ok217)* was performed with microinjection of 70 ng/µl of appropriate plasmid into already existing *syd-2(ok217)*;nthEx[*Punc-104::unc-104::mrfp*] to generate OIW 74 *syd-2(ok217)*;nthEx74[*Punc-104::unc-104::mrfp; Punc-104::gfp::syd-2*], OIW 75 *syd-2(ok217)*;nthEx75[*Punc-104::unc-104::mrfp; Punc-104::gfp::syd-2 Y741F*] and OIW 76 *syd-2(ok217)*;nthEx76[*Punc-104::unc-104::mrfp; Punc-104::gfp::syd-2 Y741E*].

Mutant strain ZM607 *syd-2(ok217)* carries a deletion of most of the N-terminal CC domains in the syd-2 gene leading to a null mutation [2]. This deletion was confirmed via PCR using the following primer set: CGCGGGAATTATGCCTATTA (forward) and TTGCATCTGCAAAAGAAACG (reverse). Mutant strain RB633 *ptp-3(ok244)* carries a deletion of three FN3 domains leading to a frameshift and premature termination of the PTP-3A isoform. The deletion was confirmed via PCR using the following primer set: ACCCAAACGTTACCGAACAG (forward) and CCAGGGACTGCAGGAAAATA (reverse). Mutant strain CZ3761 *ptp-3(mu256)* is characterized by an insertion of a single nucleotide in exon 25 of the ptp-3 gene resulting in a frameshift leading to a premature stop at AA1777 (Supp. Fig. 1) [23]. This single nucleotide insertion was confirmed by sequencing and in addition, a G to T (resulting in coding the same amino acid – leucine) mutation four nucleotides upstream of the aforementioned insertion was found. Nevertheless, the single nucleotide insertion should still result in a premature stop leading to a null mutation as described in [23]. Note that *ptp-3(mu256)* mutant worms revealed mild egg retention phenotypes (Supp. Fig. 2).

### Microinjection of *C. elegans* plasmids

Microinjection technique as described in [60] was employed with the modification that either L4 staged worms or mostly young adult worms were microinjected. Briefly, immobilized worms (on 2.5% agarose pad immersed in a small drop of mineral oil (Sigma, M5904)) were microinjected using the DIC function of an Olympus IX81 microscope fitted with a micromanipulator controlled by an Eppendorf Femtojet express. Worms were recovered using recovery buffer or S buffer and then placed onto NGM plates. F1 generations were screened for the transgenic genes and maintained separately to clone out a stable line at 20°C.

### Real-time PCR

Messenger RNA was isolated from lysates of young adult worms using TRIzol reagent (Invitrogen) following reverse transcription using the SuperScript First-Strand Synthesis system (Invitrogen). The primer set used for qPCR of the unc-104 gene was: GAAGGAAATAAAGCGAGAAC (forward) and CTGCCAAATCAACCAAAG (reverse) and the primer set for the syd-2 gene was: CGGAACAATACTCGACTTC (forward) and GCCACACGCTCCATT (reverse). Internal control for qPCR was the cdc-42 gene as described by [61] and primer set was: CTGCTGGACAGGAAGATTACG (forward) and TCGGACATTCTCGAATGAAG (reverse). ABI Power SYBR Green PCR Master Mix (ABI, USA) was used in conjunction with a PCR machine model ABI StepOne Plus Real-time PCR (Applied Biosystems). The experiment was repeated 3 times and unpaired Student’s t-test was used to compare the study groups.

### Co-immunoprecipitation assays

For co-immunoprecipitation assays (Fig. 2), 2 µg of anti-GFP (GFP antibody GT859, Gene Tex) antibody was incubated with protein G coated magnetic beads (PureProteome protein G magnetic beads, LSKMAGG10, EMD Millipore Corp.) for 1-3 hrs in IP (immunoprecipitation) buffer at 16°C. After washes, 500 µg or 1 mg of total protein extract was used to pull-down the target proteins overnight. After subsequent washes, beads were boiled in the presence of SDS-PAGE loading solution and separated on 6% SDS-PAGE. Separated proteins were then transferred to a PVDF membrane (IPVH00010, Millipore Corp.) for overnight at 4°C in a wet transfer system (Hoefer, Inc.) at 35 mA. The blot was blocked with 5% BSA and probed with primary and secondary antibodies before visualization of the band using CyECL Western blotting substrate H (ECL003-H, Cyrus bioscience) in ImageQuant LAS4000mini. Anti-GFP antibody (B-2, SC-9996, mouse monoclonal, Santa Cruz Biotechnology) was diluted at 1:10000 and anti-SYD-2 antibody (cL-19, SC-15655, goat polyclonal, Santa Cruz Biotechnology, USA) at 1:500. Secondary HRP antibody was used at a dilution of 1:10000 (anti-mouse IgG HRP secondary, GTX213111-01, Gene Tex) and 1:2500 (anti-goat IgG HRP secondary, GTX228416-01, Gene Tex), respectively. Co-IP experiments were repeated 3 times and a single representative cropped blot is presented (Fig. 1). Co-precipitated SYD-2 protein was normalized to the pulled-down UNC-104::GFP (Fig. 1). For relative quantification of co-precipitated protein levels, the normalized signals from the mutant group were once again normalized to the control group and represented as percentages (Fig. 1).

### Colocalization analysis and BiFC assays

For colocalization analysis (Fig. 2), images were collected from live worms using an inverted Zeiss LSM 780 laser scanning confocal microscope. ICQ values were then calculated from a region of interest using ImageJ plugin ‘Intensity Correlation Analysis’ after background subtraction using the plugin ‘ROI’. ICQ values range between −0.5 to +0.5, with values towards +0.5 representing two fluorophores with dependent expression patterns, and values close to −0.5 representing fluorophores that exhibit segregated expression patterns, while 0 stands for random expression. Additional to ICQ values, we present PDM images based on: Product of the Differences from the Mean = (cyan intensity – mean cyan intensity) X (green intensity – mean green intensity).

The BiFC (bimolecular fluorescence complementation) assay is a powerful tool to understand if two proteins are in close proximity for assumed functional interactions in cells. In brief, a YFP protein (Venus) is split into two halves (N-terminal Venus VN and C-terminal Venus VC) and tagged to the two test proteins. Complementation of Venus will lead to a fluorescent signal if the two test proteins are at least 7-10 nm close to each other [38, 62]. In this study we designed fusions of PTP-3B::VC155 and VN173::SYD-2 (Fig. 2).

### RNAi knockdown

RNA interference was performed by feeding worms with bacterial strains expressing dsRNA for a particular gene of interest (RNAi feeding method by [63]). RNAi feeding clones were obtained from the Julie Ahringer’s *C. elegans* RNAi feeding library (Source BioScience LifeScience, USA; provided by the *C. elegans* Core Facility, Taiwan). Feeding strains were grown overnight at 37°C on NGM plates containing ampicillin (100 µg/ml) and 1 mM IPTG. F0 worms were transferred to the respective RNAi plate and incubated at 20°C. Worms were then transferred to new plates every day and final data analysis was carried out using young adults from the cloned F1 progeny.

### Intramolecular FRET analysis

For intramolecular FRET (Förster/Fluorescence resonance energy transfer) experiments, modified versions of CFP (donor) and YFP (acceptor), such as Cypet and Ypet, were used and tagged to the N- and C-terminus of SYD-2 (FRET sensor), respectively. Cypet and Ypet have been reported to exhibit very high ratiometric signals during FRET, when compared to their traditional counterparts [64]. Plasmids encoding Cypet (pCypet-Rac1(WT), #22782) and Ypet (pYpet-PBD, #22781) sequences were taken from the Addgene repository (Addgene Inc., USA) and subcloned into worm expression plasmids as mentioned above. For intramolecular FRET, images were obtained from live worms using an inverted Zeiss LSM 780 laser scanning confocal microscope with 63X/1.40 Oil DIC M27 objective lens (shared microscope facility at Biomedical Science and Engineering Center, NTHU, Taiwan). Images were acquired at the emission wavelength of 483 nm (donor channel) and 574 nm (FRET channel) upon excitation at 405 nm for Cypet, as well as at 574 nm (acceptor channel) upon excitation at 514 nm for Ypet, respectively. Ratiometric analysis was used to understand the intramolecular folding of SYD-2 protein as described in [39, 65, 66], since the ratio of donor molecule to acceptor molecule is 1:1 in intramolecular FRET. Thus, fluorescence intensity from FRET channel (*IFRET*) was divided by the fluorescence intensity from donor channel (*ID*) to generate a ratio (*IFRET*/*ID*) for relative comparisons between different genetic backgrounds. Differences created by presumed slight changes in expression levels between individual worms (due to the extrachromosomal array nature of the FRET sensor) were normalized in the ratiometric analysis (*IFRET*/*ID*). Briefly, lower ratio value represents open conformation of FRET sensor and higher ratio value indicates intramolecularly folded conformation of FRET sensor. FRET pseudo-color images were generated by the “Fret analyzer” plugin from ImageJ that represents a 16 false color scale. FRET ratios from different study groups were assessed with one-way ANOVA using Fisher’s LSD comparison test.

### Fluorescence intensity measurements of STVs in neurons

Quantification of SNB-1::mRFP fluorescence in somas, axons, synapses, VNC (ventral nerve cord) and DNC (dorsal nerve cord) (Fig. 4) were carried out using ImageJ based on a published protocol [67]. Parameters as area, integrated density, minimum/maximum gray value and mean gray value were chosen and then the appropriate region of interest as well as the background selected and measured. The following formula was used to correct for background fluorescence: Integrated density – (area of selected region * mean gray value from the background). The total fluorescence intensity measured for a neuron was considered 100% to normalize for the variation in expression of extrachromosomal arrays between individual worms, thus, the percentage of cargo measured in subcellular regions equals to: (Fluorescence intensity in subcellular region/total fluorescent intensity in the neuron)*100. One-way ANOVA with Fisher’s LSD multiple comparison test was used to compare the study groups.

### Axonal motor and cargo cluster analysis

For quantification of UNC-104 motor clusters (Fig. 5) along sub-lateral neurons, images were obtained using an Olympus IX81 microscope connected to an Andor iXon DV897 EMCCD camera and analyzed using ImageJ. Note that particles of sizes less than 3 pixels were excluded from analysis to reject random noise. For density measurements, the number of particles along axons was manually counted and the length of the axon was noted. The following formula was used for density calculation: (Total number of particles in the neurite segment/Length of that neurite segment)*100. Analyzed particles from different groups were subjected to one-way ANOVA with multiple comparisons using Fisher’s LSD test.

For the quantification of SNB-1 particles (Fig. 4), we used an inverted Zeiss LSM 780 laser scanning confocal microscope to collect the data. The area of SNB-1 particles were analyzed as mentioned above. For analysis of distances travelled by SNB-1 cargo, a line between the axon hillock and the detected SNB-1 fluorescent particle was drawn using ImageJ’s line tool and then the measurement executed. Measured distances were normalized (distance between axon hillock and detected SNB-1 particle/total length of initial axonal region analyzed) by the total length of axon seen in the field of observation and converted into percentage of distances traveled by the particles (Fig. 4D). For particle density measurements (UNC-104 and SNB-1), the following formula was used: (Number of particles in the axon/Length of the axon)*100. One-way ANOVA with multiple comparisons using Fisher’s LSD test was used to compare the study groups.

### Motility analysis

For quantification of motility parameters of fluorescently-tagged proteins in axons of ALM neurons, worms were immobilized between cover glasses coated with 2% agarose (without using any anesthetics such as levamisole). Time-lapse imaging was carried out using an Olympus IX81 microscope fitted with a DSU Nipkow spinning disk connected to an Andor iXon DV897 EMCCD camera; image acquisition rate of 3 to 4 frames per second was used. Obtained images were then analyzed by the kymograph method using ImageJ software (https://imagej.nih.gov/ij/, NIH) [3]. Here, the “Straighten” plugin was used to straighten curved axon followed by the execution of the “Reslice stack” function to generate a kymograph. A particle pause is regarded if the velocity of a particle is less than 0.07 µm/s. Vertical lines in a kymograph indicate stationary particles, whereas horizontally angled lines correspond to moving particles. An “event” is defined as an occurrence of a movement of a fluorescent particle after a reversal or a pause. Anterograde or retrograde run length is the distance covered by the fluorescent particle in the acquired kymograph. Directional changes (either from anterograde to retrograde or from retrograde to anterograde) are defined as changes in directions of a single event. The term motility is defined as a movement occurrence of any event during the time of imaging (refer to Supp. Fig. 4 for details). Analyzed particles and obtained values from different groups were tested with one-way ANOVA with multiple comparisons using Fisher’s LSD test. Histograms were plotted (Supp. Figs. 6+7) to identify outliers.

## Data availability

Original data will be provided upon request.

## Acknowledgements

We thank Prof. Eric Hwang, National Chiao Tung University (Hsinchu, Taiwan) for his guidance in intramolecular FRET experiments. We thank Dr. Prerana Bhan for general technical guidance in our lab. We thank the *C. elegans* Core Facility (CECF) Taiwan (funded by the Ministry of Science and Technology, MOST) for providing microinjection setups and worm observation systems. We acknowledge MOST grants NSC 100-2311-B-007-004- and MOST 103-2311-B-007 - 004 -MY3 to OIW.

## Authors’ contributions

MMS – designed experiments, performed experiments, analyzed experiments and wrote the manuscript, SNB – performed and analyzed experiments, HYH – performed experiments, OIW – obtained funding, designed experiments and wrote the manuscript.

